# Incretin hormones and pharmacomimetics rapidly inhibit AgRP neuron activity to suppress appetite

**DOI:** 10.1101/2024.03.18.585583

**Authors:** Hayley E. McMorrow, Carolyn M. Lorch, Nikolas W. Hayes, Stefan W. Fleps, Joshua A. Frydman, Jessica L. Xia, Ricardo J. Samms, Lisa R. Beutler

## Abstract

Analogs of the incretin hormones glucagon-like peptide-1 (GLP-1) and glucose-dependent insulinotropic peptide (GIP) have become mainstays of obesity and diabetes management. However, both the physiologic role of incretin hormones in the control of appetite and the pharmacologic mechanisms by which incretin-mimetic drugs suppress caloric intake remain incompletely understood. Hunger-promoting AgRP-expressing neurons are an important hypothalamic population that regulates food intake. Therefore, we set out to determine how incretins analogs affect their activity *in vivo*. Using fiber photometry, we observed that both GIP receptor (GIPR) and GLP-1 receptor (GLP-1R) agonism acutely inhibit AgRP neuron activity in fasted mice and reduce the response of AgRP neurons to food. Moreover, optogenetic stimulation of AgRP neurons partially attenuated incretin-induced feeding suppression, suggesting that AgRP neuron inhibition is necessary for the full appetite-suppressing effects of incretin-based therapeutics. Finally, we found that GIP but not GLP-1 is necessary for nutrient-mediated AgRP neuron inhibition, representing a novel physiologic role for GIP in maintaining energy balance. Taken together, these findings reveal neural mechanisms underlying the efficacy of incretin-mimetic obesity therapies. Understanding these drugs’ mechanisms of action is crucial for the development of next-generation obesity pharmacotherapies with an improved therapeutic profile.

## Main

AgRP neuron activity is necessary and sufficient to promote food intake and critical for maintaining energy homeostasis^1–5^. These neurons integrate external sensory stimuli and interoceptive signals from the gastrointestinal tract to promote adaptive feeding behavior^6–10^. Through incompletely understood mechanisms, AgRP neuron inhibition by recently consumed nutrients reduces subsequent food intake via multiple gut-derived signals and neural circuits^7,11–15^. Here, we set out to investigate the effects of incretin hormones on AgRP neuron activity.

We equipped mice for *in vivo* imaging of AgRP neurons using fiber photometry. Intraperitoneal (IP) injection of the GIP analog (D-Ala^2^)-GIP (DA-GIP) or the GLP-1 analog Exendin-4 (Ex-4) rapidly inhibited AgRP neuron activity (**Fig 1**). This is consistent with *ex vivo* studies showing that GLP-1 analogs inhibit AgRP neurons^16,17^. The response of AgRP neurons to incretin analogs was dose-dependent (**Fig S1A–H, S2A–H**). Ex-4 induced greater neural inhibition than DA-GIP, and AgRP neuron inhibition in response to the combination of Ex-4 and DA-GIP was stronger than the response to Ex-4 alone (**Fig. 1A–E**).

**Figure 1.**
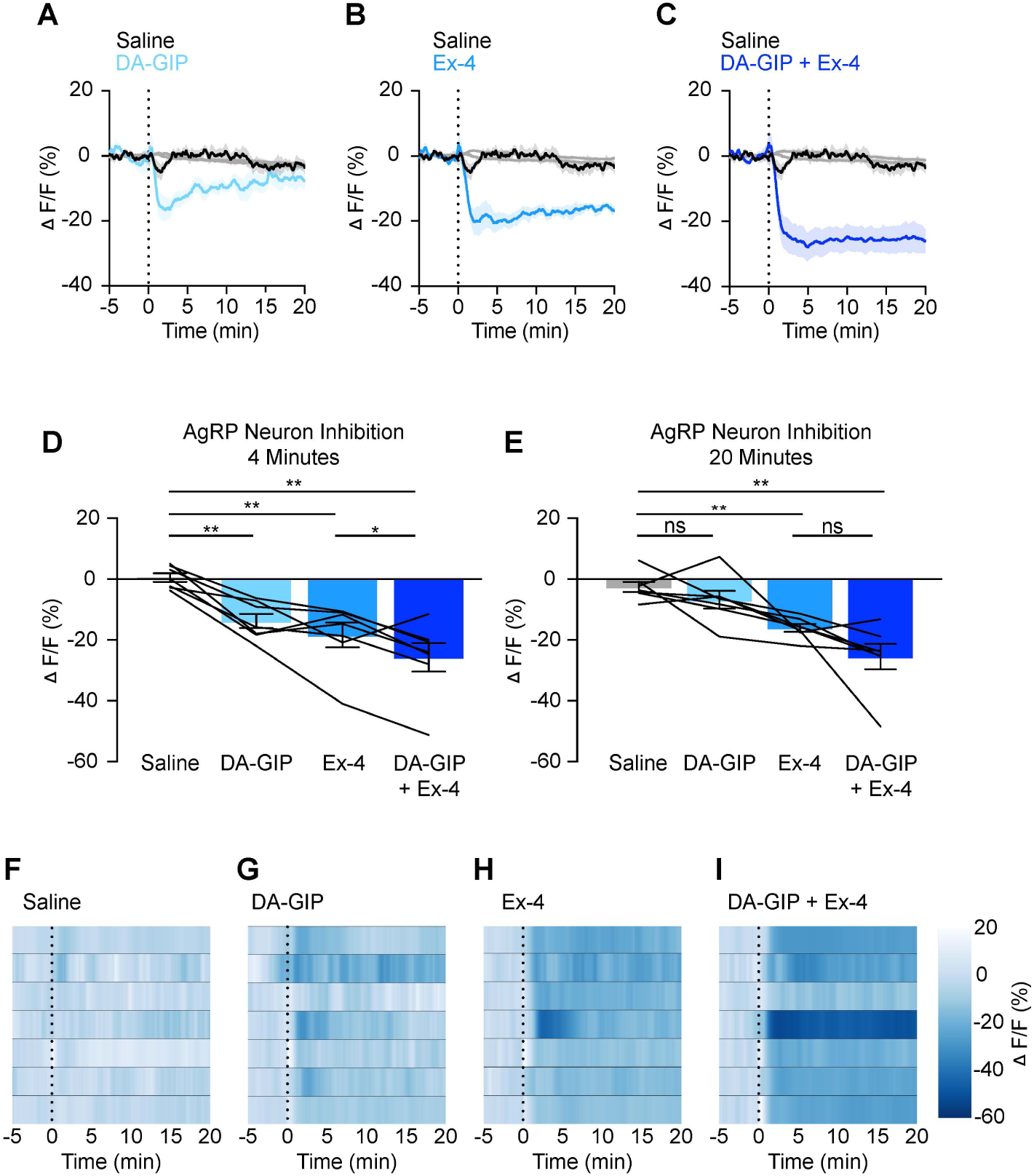
GIPR and GLP-1R agonists acutely inhibit AgRP neurons. **(A-C)** Calcium signal in AgRP neurons from fasted mice injected with DA-GIP (A), Ex-4 (B), or DA-GIP and Ex-4 (C) compared to saline as indicated. n = 7 mice per group. **(D,E)** Average ΔF/F in mice from (A-C) 4 minutes (D) and 20 minutes (E) after injection. ((D) one-way ANOVA, p<0.0001; (E) one-way ANOVA, p=0.0003). **(F-I)** Heat maps showing ΔF/F in individual mice injected with saline (F), DA-GIP (G), Ex-4 (H), or DA-GIP and Ex-4 (I). (A-C) Isosbestic traces for all recordings are shown in gray. (A-C, F-I) Vertical dashed lines indicate the time of injection. (D,E) Lines represent individual mice. Error bars indicate mean ± SEM. Post-hoc comparisons: *p<0.05, **p<0.01.

In addition to post-ingestive feedback, chow presentation induces rapid, pre-consummatory AgRP neuron inhibition with a magnitude of inhibition that correlates with the quantity of subsequent food intake^6–10^. As previously shown for the long-acting GLP-1R agonist liraglutide^16^, pre-treatment with DA-GIP or Ex-4 blunted subsequent chow-induced AgRP neuron inhibition compared to pretreatment with saline in the same mice (**Fig. 2A–H**). Remarkably, when given in combination, Ex-4 and DA-GIP suppressed chow-induced neuron inhibition more than Ex-4 alone (**Fig. 2C, D**). Reduced AgRP neuron responses to chow presentation correlated with feeding suppression induced by DA-GIP, Ex-4 or, DA-GIP + Ex-4 in fasted wildtype mice (**Fig. 2I**). Specifically, consistent with prior findings, acute treatment with DA-GIP modestly inhibited fast re-feeding but significantly potentiated the suppression of food intake induced by Ex-4^18^. The effect of Ex-4 on chow-induced AgRP neuron inhibition was dose-dependent (**Fig. S2I–P**), consistent with dose-dependent effects of GLP-1 analogs on food intake^19^. By contrast, the effect of DA-GIP on chow-induced AgRP neuron inhibition did not vary significantly with dose (**Fig. S1I–P)**, in line with the more subtle acute effects of DA-GIP on food intake. Taken together, the additive effect of GIPR and GLP-1R agonism on AgRP neuron dynamics aligns with mounting evidence for the superior efficacy of combined GIP and GLP-1R activation when compared to selective GLP-1R monoagonism for the treatment of obesity^20–24^ .

**Figure 2.**
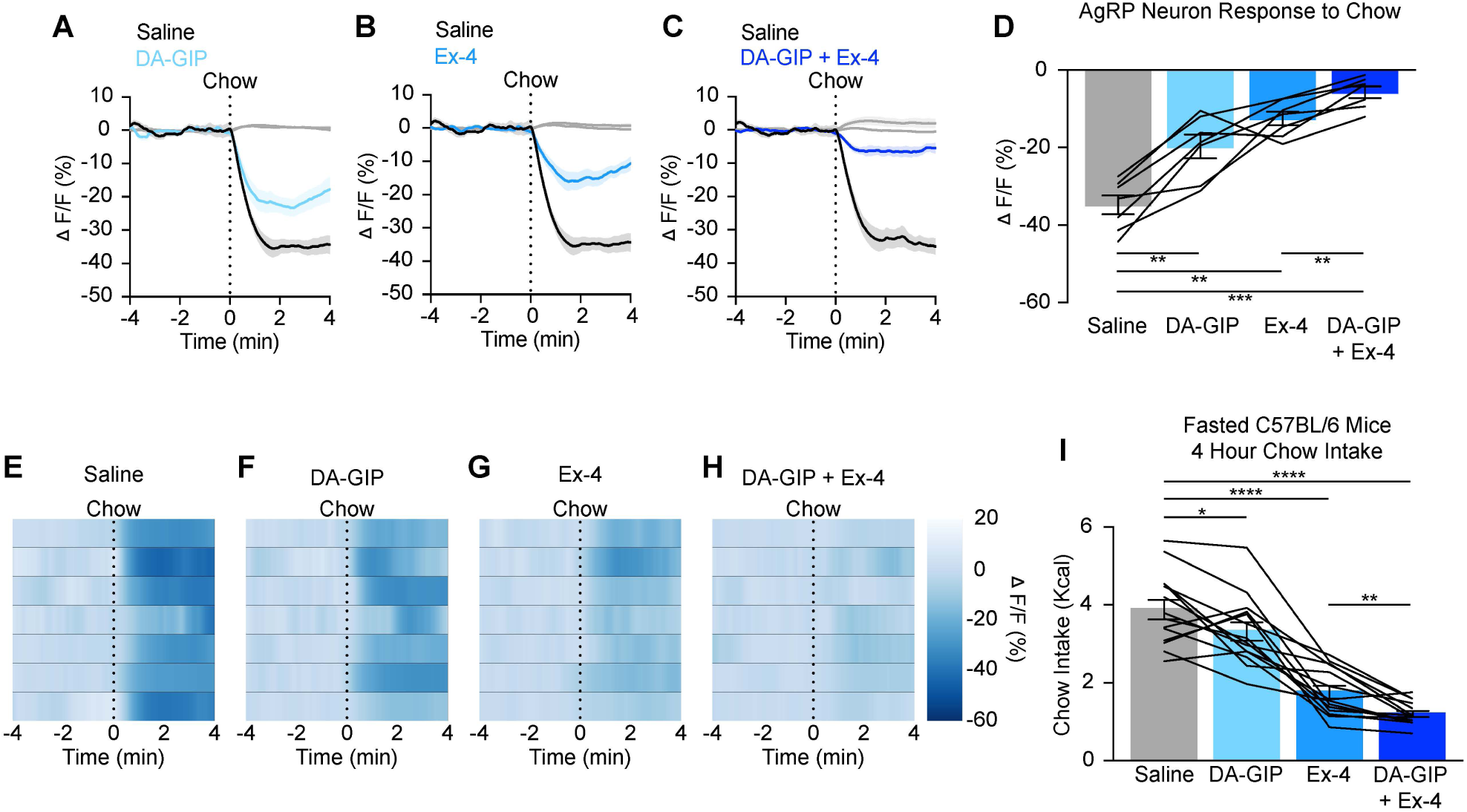
GIPR and GLP-1R agonists additively attenuate the AgRP neuron response to food presentation and food intake. **(A-C)** Calcium signal in AgRP neurons from fasted mice presented with chow 20 minutes after pre-treatment with DA-GIP (A), Ex-4 (B), or DA-GIP and Ex-4 (C) compared to saline as indicated. n = 7 mice per group. **(D)** Average ΔF/F in mice from (A-C) 4 minutes after chow presentation. (one-way ANOVA, p=<0.0001). **(E-H)** Heat maps showing ΔF/F in individual mice from (A-C) after chow presentation. **(I)** Four-hour chow intake following a five-hour fast and incretin or saline injection as indicated in C57BL/6 mice. n = 14 mice per group. (one-way ANOVA, p =<0.0001). (A-C) Isosbestic traces for all recordings are shown in gray. (A-C, E-H) Vertical dashed lines indicate the time of chow presentation. (D,I) Lines represent individual mice. Error bars indicate mean ± SEM. Post-hoc comparisons: *p<0.05, **p<0.01, ***p<0.001, ****p<0.0001.

We next used an optogenetic approach to investigate the behavioral relevance of incretin-mediated AgRP neuron inhibition. To determine whether AgRP neuron stimulation can overcome incretin receptor agonist-induced feeding suppression, mice that express channelrhodopsin2 (ChR2) selectively in AgRP neurons (AgRP::ChR2 mice) were equipped for optogenetic stimulation of AgRP neuron cell bodies. These mice were fasted for five hours, habituated to feeding chambers for 30 minutes, then systemically treated with saline, Ex-4, or Ex-4 + DA-GIP and immediately re-fed in the absence or presence of light stimulation (**Fig. 3A**). In saline-treated mice, AgRP neuron stimulation significantly increased food intake as expected (**Fig. 3B**). AgRP neuron stimulation partially rescued the anorexia induced by both Ex-4 and Ex-4 + DA-GIP (**Fig. 3B**). Thus, AgRP neuron inhibition likely contributes to incretin analog-induced appetite suppression.

**Figure 3.**
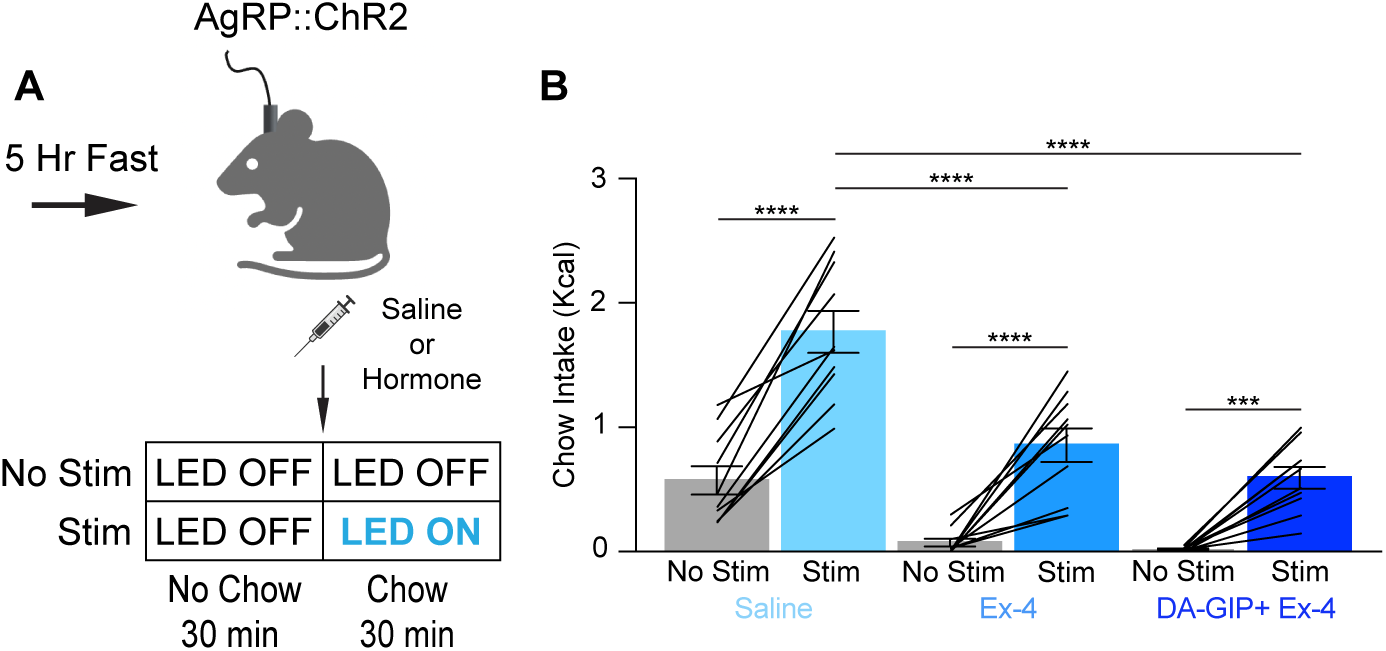
AgRP neuron stimulation partially rescues acute incretin-induced feeding suppression. **(A)** Experimental schematic. **(B)** 30-minute chow intake in fasted mice following injection of saline, Ex-4, or DA-GIP and Ex-4 in the presence or absence of AgRP neuron stimulation as indicated. n = 10 mice per group. (two-way ANOVA, main effect of hormone treatment p=<0.0001, main effect of no stim vs. stim p=<0.0001, interaction, p=0.0036). Lines represent individual mice. Error bars indicate mean ± SEM. Post-hoc comparisons: ***p<0.001, ****p<0.0001.

These experiments focused on the pharmacologic effects of incretin hormones on AgRP neurons. The physiologic effects of incretins on hypothalamic feeding circuits are also incompletely understood. Specifically, it is unclear whether GIPR and/or GLP-1R signaling are necessary for nutrient-mediated AgRP neuron inhibition^7,8^. To examine this, mice were equipped for fiber photometry recording from AgRP neurons and intragastric infusion of nutrients^7^, and neural responses to nutrients were measured in the presence versus absence of incretin receptor blockade. To examine the role of GIPR, we first pre-treated mice with a control (non-neutralizing) antibody, then intragastrically administered glucose, lipid, or Ensure on different days. Following intragastric nutrient infusions under control conditions, mice were treated with a long-acting, neutralizing monoclonal murine GIPR blocking antibody (muGIPR-Ab)^25^, and nutrient infusions were repeated. GIPR blockade attenuated glucose- and Ensure-mediated AgRP neuron inhibition, but not lipid-induced AgRP neuron inhibition (**Fig. 4**). Because muGIPR-Ab is long-acting, the order of pre-treatments could not be counterbalanced. However, control experiments showed that mice maintained consistent neural responses to repeated intragastric nutrient infusions over one to two weeks in the absence of antibody treatment (**Fig. S3**), and multiple prior studies have shown consistent nutrient-mediated AgRP neural responses for several weeks in control mice^26–28^. In contrast to the dramatic effects of GIPR blockade, pretreatment with the GLP-1R antagonist exendin 9-39 (Ex-9) had no effect on nutrient-mediated AgRP neuron inhibition (**Fig. S4**). Finally, neither the GIPR blocking antibody nor Ex-9 dramatically impacted AgRP neuron inhibition in response to food presentation (**Fig. S5 A–G**). This suggests that the blunted responses to gastrointestinal nutrients following muGIPR-Ab are not likely due to a floor effect in the setting of altered baseline AgRP neuron activity. The very small but statistically significant effect of muGIPR-Ab on chow-mediated AgRP neuron inhibition we observed may be related to its previously reported modest effect on food intake (**Fig S5 A, B**)^25,29^. This newly described physiological function for GIP in mediating nutrient-dependent AgRP neuron inhibition may partially underlie the enhanced weight loss efficacy of dual GIP and GLP-1R agonists when compared to GLP-1R monoagonists, as GIPR activation may recapitulate the post-ingestive effects of glucose.

**Figure 4.**
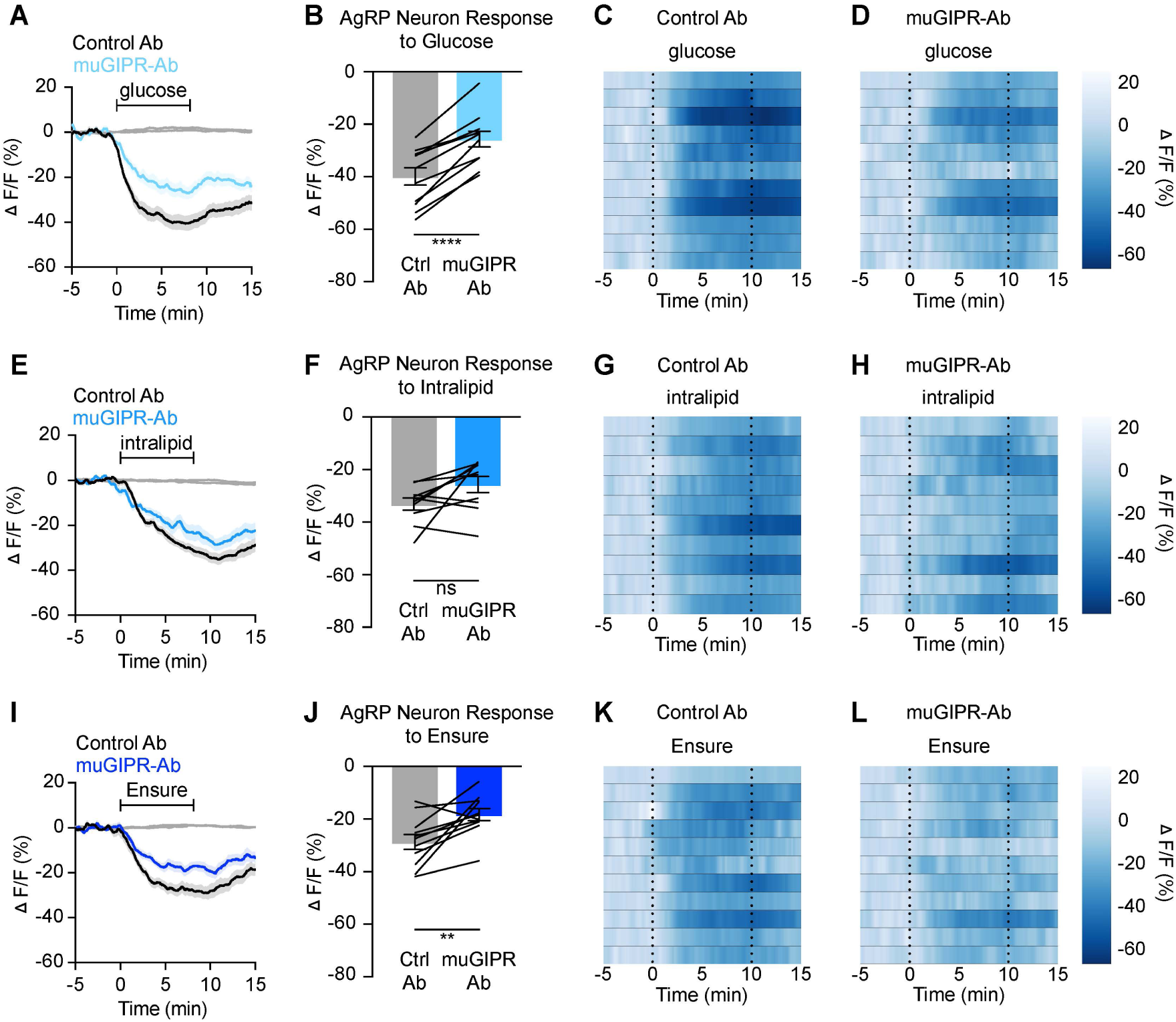
Signaling through GIPR is necessary for nutrient-mediated AgRP neuron inhibition. **(A,E,I)** Calcium signal in AgRP neurons from fasted mice during infusion of glucose (A), intralipid (E), or Ensure (I) after pre-treatment with control or muGIPR-Ab as indicated. n = 10-11 mice per group. **(B,F,J)** Average ΔF/F in mice from (A,E,I) at the end of nutrient infusion. ((B) paired t-test, p=<0.0001; (F) paired t-test, p=0.0557; (J) paired t-test, p=0.0061). **(C,D,G,H,K,L)** Heat maps showing ΔF/F in individual mice from (A,E,I) during nutrient infusion. (A,E,I) Isosbestic traces for all recordings are shown in gray. (C,D,G,H,K,L) Vertical dashed lines indicate the start and end of nutrient infusions. (B,F,J) Lines represent individual mice. Error bars indicate mean ± SEM. T-tests: **p<0.01, ****p<0.0001.

Taken together, these findings reveal novel roles for AgRP neurons in incretin pharmacology and physiology. In particular, the physiologic function of GIP in food intake and body weight maintenance has remained elusive. Numerous studies have shown that GIPR agonism reduces food intake^30,31^. By contrast, other studies have shown that global GIPR knockout mice are protected from both obesity and insulin resistance when fed a high-fat diet^32–36^, and GIPR antagonism coupled with GLP-1R agonism leads to weight loss in early clinical trials^37,38^ and mouse models^25,29^. Here, we have identified a clear role for GIP in gut-brain communication with AgRP neurons in a manner that is consistent with the anorexigenic effects of GIPR agonism. Further studies will be required to define which GIPR-expressing neurons are required to elicit this effect, and how obesity impacts gut-brain axis responsiveness to incretin hormones *in vivo*^26–28^.

In addition to illuminating a previously unknown physiologic role of GIP, we have also shown that AgRP neurons play a critical role in mediating the anorexigenic effects of pharmacologic incretin receptor agonism. GLP-1 and GIP analogs rapidly inhibit AgRP neurons, and stimulation of AgRP neurons partially restores food intake following treatment with these incretin agonists, suggesting that AgRP neuron inhibition contributes to the anorexigenic effect of incretin-mimetic therapies. While these findings add significantly to our mechanistic understanding of incretin-based anti-obesity agents, many questions remain to be addressed.

It is unclear what cell types and circuits GLP-1R and GIPR agonists act upon to inhibit AgRP neurons, though based on prior *ex vivo* physiology studies and RNA sequencing findings, this effect is likely indirect^16,17,39–41^. The GLP-1R is expressed in feeding-related nuclei in the hypothalamus and brainstem^42,43^, and knockout from glutamatergic but not GABAergic neurons almost entirely abrogates GLP-1R agonist-induced weight loss in obese mice^44^. While hypothalamic or hindbrain knockdown of GLP-1R reduces liraglutide efficacy, no brain region has been shown to be solely responsible for the weight loss efficacy of this drug^17,45,46^. A recent study described a population of GLP-1R agonist-activated neurons in the arcuate nucleus that inhibit AgRP neurons; however, whether direct GLP-1R activation of this cell type contributes to incretin-mimetic induced weight loss has yet to be illuminated^47^. The GLP-1R is also expressed in a large population of vagal afferent neurons^48–51^, and chemogenetic activation of these distension-sensing nodose ganglion neurons is sufficient to inhibit AgRP neurons^50^. Moreover, central blockade of the GLP-1R does not abrogate the anorexic effects of peripherally administered Ex-4^52^, and GLP-1R deletion from peripheral sensory neurons modestly attenuates the appetite suppressing and weight loss efficacy of GLP-1R agonists in obese mice^53,54^. Thus, GLP-1-induced appetite suppression and weight loss may be mediated by multiple peripheral and central neural circuits^55^. Alongside prior studies, our data suggest that AgRP neurons are an indirect but critical target of GLP-1-based therapies.

Similarly, the GIPR is expressed in hypothalamic feeding centers, area postrema and NTS but not in hypothalamic AgRP neurons^40,41^. CNS knockout of the GIPR from GABAergic neurons blocks the modest anorectic effects of long-acting GIPR agonists and abrogates the benefit of dual GLP-1 and GIP receptor agonism when compared to GLP-1R agonism in obese mice^31,40,56^. Chemogenetic activation of GIPR-expressing cells in the hypothalamus or dorsal vagal complex reduces feeding, but local GIPR knockout in the hypothalamus does not blunt incretin-mimetic induced weight loss^40,57^. The GIPR is also expressed at low levels in nodose and dorsal root ganglia, but its function in these cell populations has not been examined^48,57–59^. Recent data support a critical role for spinal afferent neurons in glucose-mediated AgRP neuron inhibition^12^, and it is possible that GIPR activation in peripheral sensory neurons is required for this. Additional studies examining the effect on feeding and neural activity of local, cell-type specific GIPR knockout will be necessary to clarify the physiologic and pharmacologic roles of this hormone.

In summary, gut hormone receptor agonism has ushered in a new era of obesity management with the efficacy of multi-receptor agonism rivaling that of bariatric surgery. Understanding the molecular and circuit-based mechanisms of hormone-mediated appetite control is critical to refine and more precisely target future therapies to the key cell types mediating the transformative effects of these drugs. Using modern neuroscience and genetic approaches, we have dissected the role of AgRP neurons in incretin-mediated gut-brain communication and elucidated previously unreported physiologic and pharmacologic effects of GLP-1 and GIP on this axis.

## Acknowledgments

We thank Dr. Joseph Bass and Dr. Jones G. Parker for providing feedback on the manuscript. This work was supported by the American Diabetes Association Pathway to Stop Diabetes Award (12-22-ACE-31) and by NIH grants (P30-DK020595), (K08-DK118188), and (R01-DK128477) (L.R.B.).

## Figure Titles and Legends

**Figure S1.**
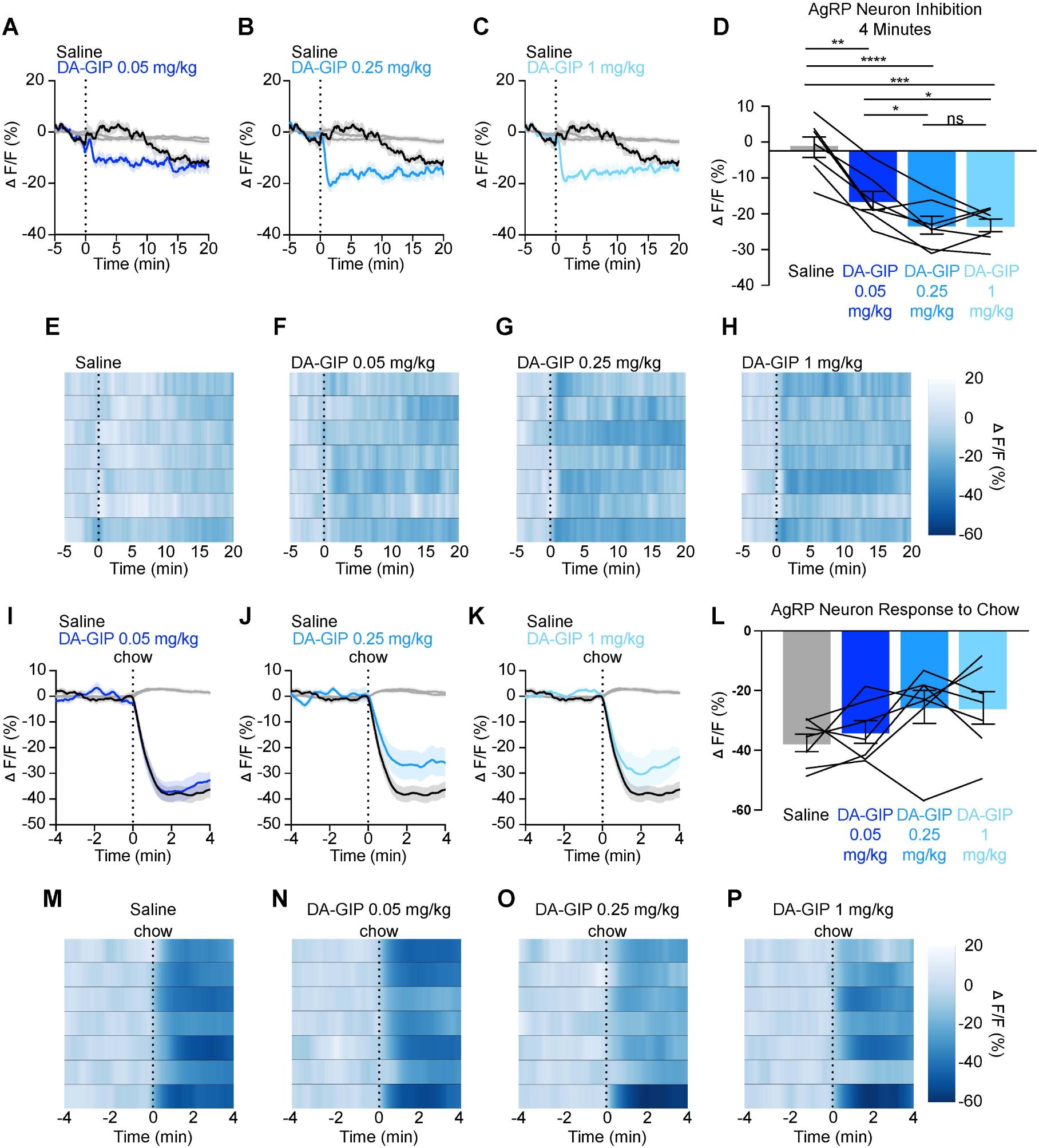
AgRP neuron responses to GIPR agonists are dose-dependent. **(A-C)** Calcium signal in AgRP neurons from fasted mice injected with DA-GIP at 0.05 mg/kg (A), 0.25 mg/kg (B), or 1 mg/kg (C) compared to saline as indicated. n = 7 mice per group. **(D)** Average ΔF/F in mice from (A-C) 4 minutes after injection. (one-way ANOVA, p=<0.0001). **(E-H)** Heat maps showing ΔF/F in individual mice from (A-C) injected with saline (E), DA-GIP at 0.05 mg/kg (F), 0.25 mg/kg (G), or 1 mg/kg (H) as indicated. **(I-K)** Calcium signal in AgRP neurons from fasted mice presented with chow 20 minutes after pre-treatment with DA-GIP at 0.05 mg/kg (I), 0.25 mg/kg (J), or 1 mg/kg (K) compared to saline as indicated. n = 7 mice per group. **(L)** Average ΔF/F in mice from (I-K) 4 minutes after chow presentation. (one-way ANOVA, p=0.0738). **(M-P)** Heat maps showing ΔF/F in individual mice from (I-K) after chow presentation. (A-C, I-K) Isosbestic traces for all recordings are shown in gray. (A-C, E-H, I-K, M-P) Vertical dashed lines indicate the time of injection or chow presentation. (D,L) Lines represent individual mice. Error bars indicate mean ± SEM. Post-hoc comparisons: *p<0.05, **p<0.01, ***p<0.001, ****p<0.0001.

**Figure S2.**
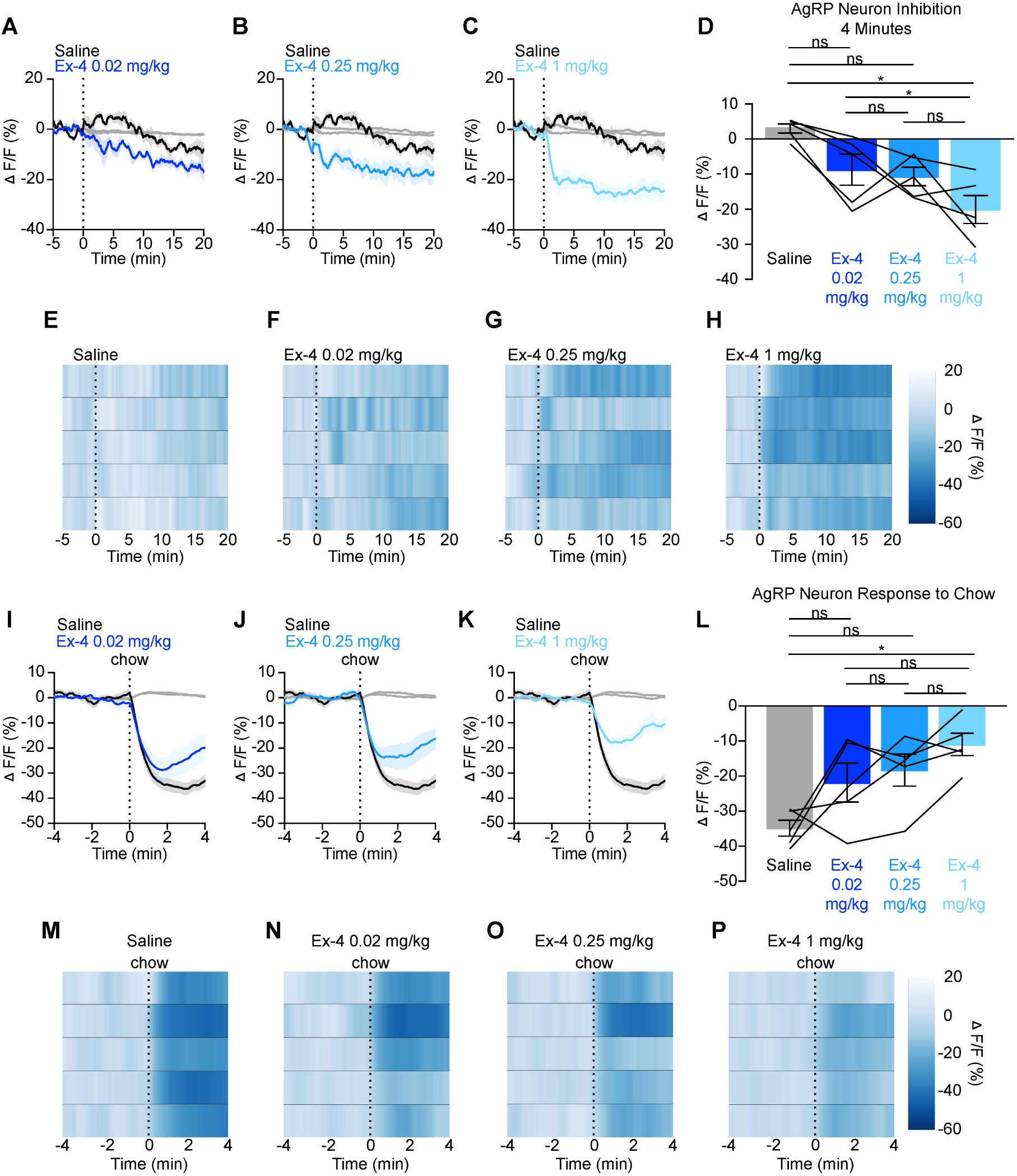
AgRP neuron responses to GLP-1R agonists are dose-dependent. **(A-C)** Calcium signal in AgRP neurons from fasted mice injected with Ex-4 at 0.02 mg/kg (A), 0.25 mg/kg (B), or 1 mg/kg (C) compared to saline as indicated. n = 5 mice per group. **(D)** Average ΔF/F in mice from (A-C) 4 minutes after injection. (one-way ANOVA, p=0.0119). **(E-H)** Heat maps showing ΔF/F in individual mice from (A-C) injected with saline (E), Ex-4 at 0.02 mg/kg (F), 0.25 mg/kg (G), or 1 mg/kg (H) as indicated. **(I-K)** Calcium signal in AgRP neurons from fasted mice presented with chow 20 minutes after pre-treatment with Ex-4 at 0.02 mg/kg (I), 0.25 mg/kg (J), or 1 mg/kg (K) compared to saline as indicated. n = 5 mice per group. **(L)** Average ΔF/F in mice from (I-K) 4 minutes after chow presentation. (one-way ANOVA, p=0.0219). **(M-P)** Heat maps showing ΔF/F in individual mice from (I-K) after chow presentation. (A-C, I-K) Isosbestic traces for all recordings are shown in gray. (A-C, E-H, I-K, M-P) Vertical dashed lines indicate the time of injection or chow presentation. (D,L) Lines represent individual mice. Error bars indicate mean ± SEM. Post-hoc comparisons: *p<0.05.

**Figure S3.**
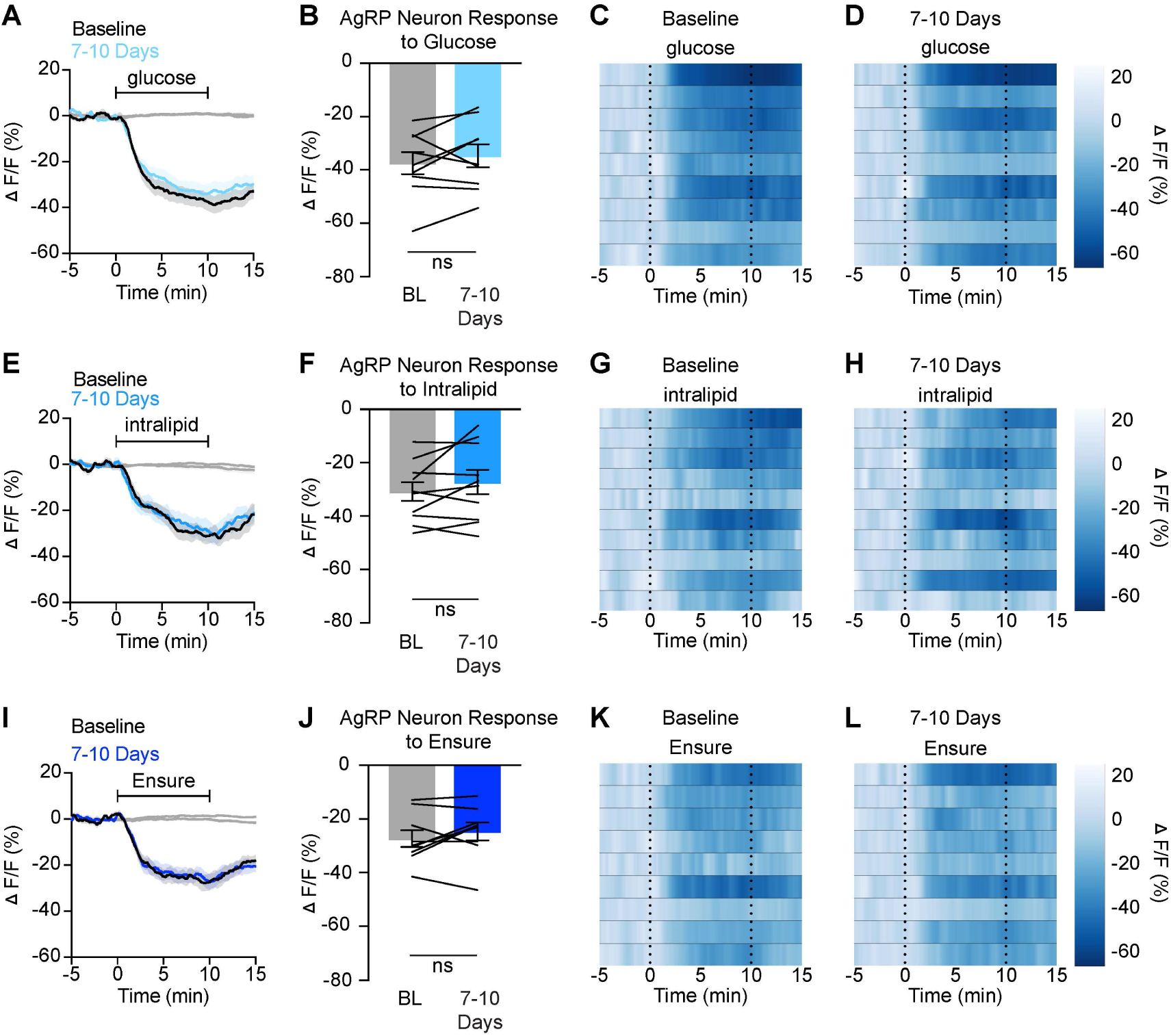
Nutrient-mediated AgRP neuron inhibition is stable over time in untreated mice. **(A,E,I)** Calcium signal in AgRP neurons from fasted mice during infusion of glucose (A), intralipid (E), and Ensure (I) at baseline and 7-10 days later as indicated. n = 9-10 mice per group. **(B,F,J)** Average ΔF/F in mice from (A,E,I) at the end of nutrient infusion. ((B) paired t-test, p=0.3471; (F) paired t-test, p=0.1781; (J) paired t-test, p=0.2725). **(C,D,G,H,K,L)** Heat maps showing ΔF/F in individual mice from (A,E,I) during nutrient infusion. (A,E,I) Isosbestic traces for all recordings are shown in gray. (C,D,G,H,K,L) Vertical dashed lines indicate the start and end of nutrient infusions. (B F,J) Lines represent individual mice. Error bars indicate mean ± SEM.

**Figure S4.**
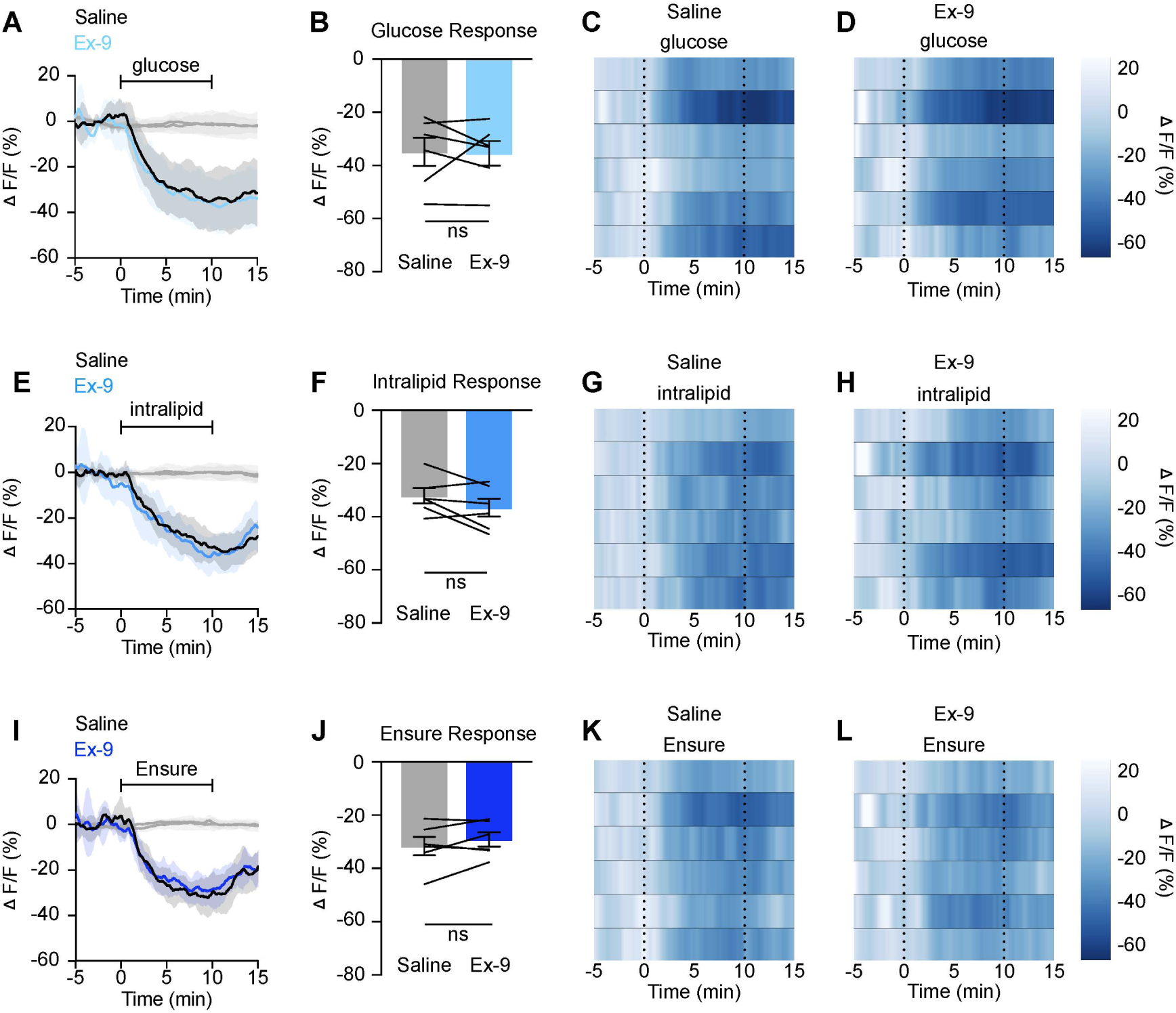
Signaling through GLP-1R is not necessary for nutrient-mediated AgRP neuron inhibition. **(A,E,I)** Calcium signal in AgRP neurons from fasted mice during infusion of glucose (A), intralipid (E), or Ensure (I) after pre-treatment with saline or Ex-9 as indicated. n = 6 mice per group. **(B,F,J)** Average ΔF/F in mice from (A,E,I) at the end of nutrient infusion. ((B) paired t-test, p=0.8918; (F) paired t-test, p=0.1314; (J) paired t-test, p=0.2401). **(C,D,G,H,K,L)** Heat maps showing ΔF/F in individual mice from (A,E,I) during nutrient infusion. (A,E,I) Isosbestic traces for all recordings are shown in gray. (C,D,G,H,K,L) Vertical dashed lines indicate the start and end of nutrient infusions. (B,F,J) Lines represent individual mice. Error bars indicate mean ± SEM.

**Figure S5.**
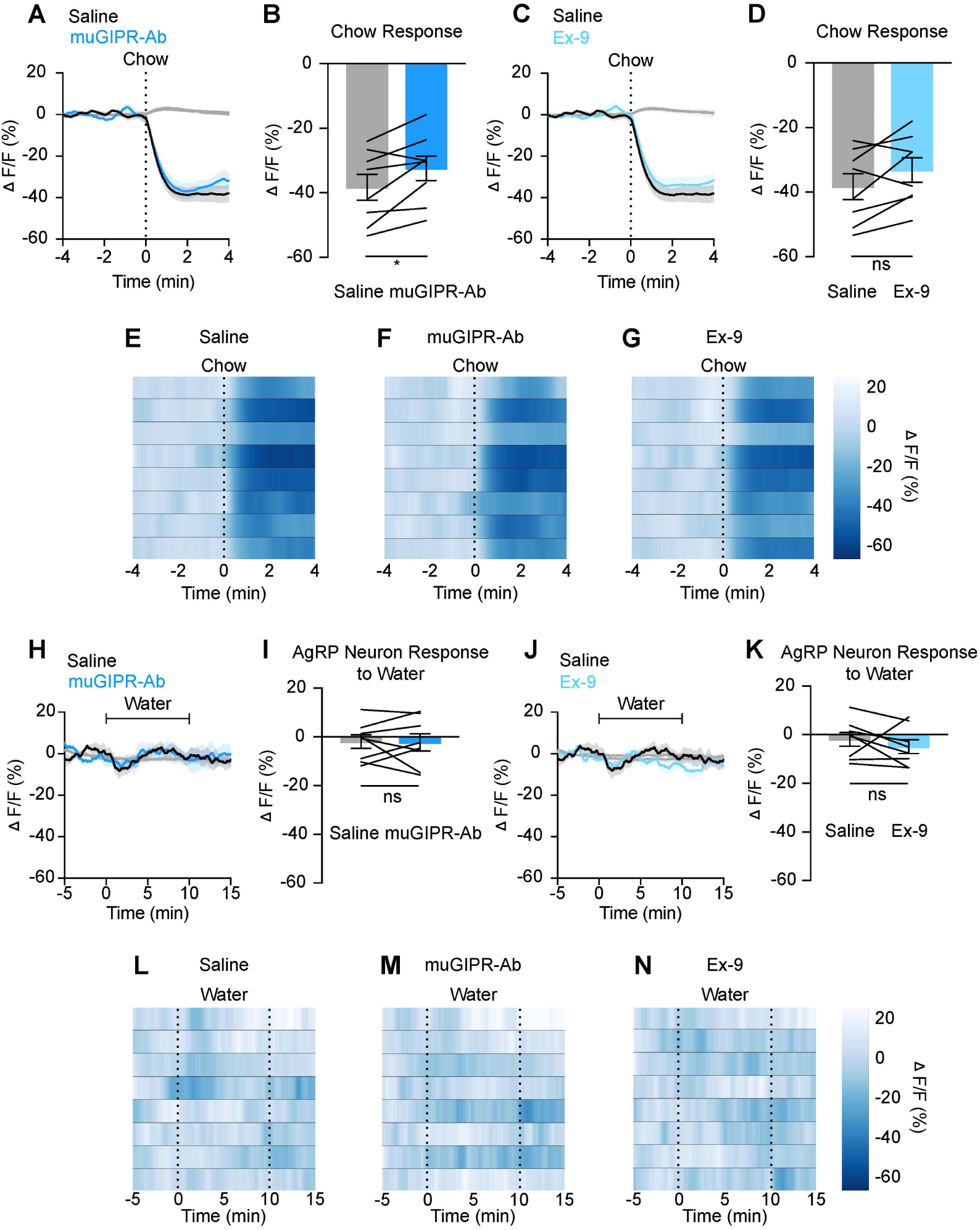
GIPR and GLP-1R blockade minimally impact AgRP neural response to food presentation or water infusion. **(A,C)** Calcium signal in AgRP neurons from fasted mice presented with chow after pre-treatment with muGIPR-Ab (A), Ex-9 (C) or saline as indicated. n = 8 mice per group. **(B,D)** Average ΔF/F in mice from (A,C) 4 minutes after chow presentation. ((B) paired t-test, p=0.0258; (D) paired t-test, p=0.0792). **(E,F,G)** Heat maps showing ΔF/F in individual mice from (A,C) after chow presentation. **(H,J)** Calcium signal in AgRP neurons from fasted mice during water infusion after pre-treatment with muGIPR-Ab (H), Ex-9 (J) or saline as indicated. n = 8 mice per group. **(I,K)** Average ΔF/F in mice from (H,J) at the end of water infusion. ((I) paired t-test, p=0.9004; (K) paired t-test, p=0.3644). **(L,M,N)** Heat maps showing ΔF/F in individual mice from (H,J) during water infusion. (A,C,H,J) Isosbestic traces for all recordings are shown in gray. (A,C,E,F,G,L,M,N) Vertical dashed lines indicate chow presentation or the start and end of water infusions. (B,D,I,K) Lines represent individual mice. Error bars indicate mean ± SEM. T-tests: *p<0.05.

## Methods

### Animals

Experimental protocols were approved by the Northwestern University IACUC in accordance with NIH guidelines for the Care and Use of Laboratory Animals. Mice were housed in a 12/12-hour reverse light/dark cycle with *ad libitum* chow (Envigo, 7012, Teklad LM-495 Mouse/Rat Sterilizable Diet) and water access. Mice were fasted for 5 or 16 hours before experiments, as indicated in the text and figures. During fasting periods, mice had *ad libitum* water access. *Agrp*^tm1(cre)Lowl^ (AgRP-Cre, #012899, Jackson Labs) animals backcrossed onto a C57BL/6J background were used for fiber photometry and nutrient infusion experiments. For optogenetic experiments, AgRP-Cre mice were crossed with B6.Cg-*Gt(ROSA)26Sort*^m32(CAG– COP4*H124R/EYFP)Hze^/J mice (ROSA26-loxStoplox-ChR2-eYFP, #024109, Jackson Labs), to generate AgRP::ChR2 animals. C57BL/6J mice (wildtype #000664, Jackson Labs) were used to measure food intake following hormone injections. Experiments were performed in male and female mice 2-6 months of age unless otherwise indicated. Male and female data were combined. Experiments were performed during the dark cycle in a dark environment.

### Stereotaxic Surgery

For photometry experiments, AAV expressing Cre-dependent GCaMP6s (100842-AAV9, AAV9.CAG.Flex.GCaMP6s, Addgene) was injected unilaterally above the arcuate nucleus (ARC) of AgRP-Cre mice. During the same surgery, an optical fiber (MFC_400/430-0.48_6.3mm_MF2.5_FLT, Doric Lenses) was implanted unilaterally at the coordinates x = +0.25 mm, y = -1.65 mm, z = -5.95 mm from bregma. Mice were allowed 2 weeks for recovery and viral expression before beginning experiments or implanting intragastric catheters.

For optogenetic experiments, fiberoptic implants (MFC_200/245_0.37_6.1mm_ZF1.25_FLT, Doric Lenses) were placed unilaterally above the ARC of AgRP::ChR2 mice at the coordinates x = +0.25 mm, y = -1.63 mm, z = -5.85 mm from bregma. Mice were allowed 10 days for recovery during which they were habituated to handling, intraperitoneal injection, and tethering to patch cords in feeding chambers before performing experiments.

Following both surgeries, mice were treated with meloxicam and buprenorphine.

### Intragastric Catheter Implantation

Surgery was performed as previously described^7,60^. AgRP-Cre mice with working photometry implants were anesthetized with ketamine/xylazine. An incision was made between the scapula, and the skin was dissected from the subcutaneous tissue. An abdominal incision was made from the xyphoid process caudally. A sterilized catheter was pulled into the abdominal cavity via a small puncture in the abdominal wall. The stomach was externalized, punctured, and the catheter was inserted into the puncture site and sutured in place. The stomach was returned to the abdominal cavity and the abdominal muscle and skin were sutured. Lastly, the catheter was secured at its intrascapular cite using a felt button (VABM1B/22, Instech Laboratories), and the intrascapular skin incision was sutured. Post-operatively, mice were treated with meloxicam, buprenorphine, and a dose of enrofloxacin, and allowed 14 days to recover before experiments.

### Fiber Photometry

Two photometry processors were used in this study (RZ5P and RZ10X, TDT). For the RZ5P setup, the LEDs and LED driver are separate from the processor (DC4100 (LED driver); M405FP1 and M470F3 (LEDs), Thorlabs), while the RZ10X processor has these components integrated. Each mouse was run on the same system using the same patch cord for every recording session to allow for reliable within-mouse comparisons.

Blue LED (465-470 nm) and UV LED (405 nm) were used as excitation light sources. LEDs were modulated at distinct rates and delivered to a fluorescence minicube (Doric Lenses) before connecting to the mouse implants (MFC_400/430-0.48_6.3mm_MF2.5_FLT, Doric Lenses) via patch cords (MFP_400/430/1100-0.57_2m_FCM-MF2.5_LAF, Doric Lenses). Emissions were collected through the patch cords to photoreceivers (Newport Visible Femtowatt Photoreceiver for the RZ5P system; integrated Lux photosensors in the RZ10X system). Digital signals were demodulated, lock-in amplified, and collected through the processors. Data were collected using Synapse software (TDT).

During recordings, mice were placed in operant chambers (ENV-307W-CT, Med Associates) within light- and sound-attenuating cubicles (ENV-022MD, Med Associates) with no food or water access unless otherwise indicated. Mice with AgRP signals inhibited less than 20% by chow presentation were considered technical failures and excluded from further experiments.

### Hormone Injections

Exendin-4 (Ex-4) (HY-13443, MedChemExpress) and (D-Ala^2^)-GIP (DA-GIP) (4054476, Bachem) were injected intraperitoneally (IP) at the doses indicated in the text and figure legends. Where indicated, Ex-4 and DA-GIP were diluted in the same solution and injected simultaneously. All hormones were dissolved in saline. To monitor AgRP neural response to hormone treatment using fiber photometry, AgRP-Cre mice were habituated to handling, photometry recording chambers and IP injections. For recordings, mice were placed in the chambers for 20 minutes prior to injection. Following injection, the photometry recording continued for 20 minutes. To evaluate the effects of hormones on the response of AgRP neurons to food presentation, we presented mice with chow 20 minutes after hormone injection. Recording continued for 4 minutes following chow presentation.

To evaluate the effects of Ex-4 and DA-GIP on food intake, wildtype C57BL/6J mice were habituated to handling, IP injection, and individual feeding chambers before undergoing a 5 hour fast at the start of dark cycle. Following the fast, mice received an IP injection of saline, DA-GIP (1 mg/kg), Ex-4 (0.02 mg/kg) or DA-GIP and Ex-4 given simultaneously. Mice were immediately re-fed in feeding chambers and food consumption was measured at 4 hours. Each mouse received all treatments on different days and treatment order was counterbalanced.

### Optogenetic Feeding Experiments

AgRP::ChR2 mice were group-housed and ranged from 4 to 12 months old. For 10 days during recovery from surgery, mice were habituated to handling, recording chambers, and patch cord tethering. An LED source and TTL pulse generator (D-OG-LED-B/B, Prizmatix) were used to generate blue light (460 nm, 2 s ON/3 s OFF, 10 ms pulse width, 20 Hz, 10-20 mW at the fiber tip). Fiber optic patch cables (500um POF N.A. 0.63 L=75cm, Prizmatix) were connected to the mouse implants (MFC_200/245-0.37_6.1mm_ZF1.25_FLT, Doric Lenses) via a sleeve (MFC_200/245-0.37_6.1mm_ZF1.25_FLT, Doric Lenses).

On test days, mice were given 30 minutes of habituation without LED stimulation or chow. Following habituation, mice received an IP injection of saline, Ex-4 (0.02 mg/kg), or DA-GIP (1 mg/kg) plus Ex-4 simultaneously. After injection, mice were immediately given 30 minutes of access to chow with or without light stimulation. Each experiment was performed in the fasted state (5 hours, beginning at start of dark cycle) in the same mice on different days. Hormone treatment order was counterbalanced.

### Nutrient Infusions during fiber photometry recording

Nutrients were infused via intragastric catheters using a syringe pump during fiber photometry recordings as previously described^7^. All infusions were given at 0.1 mL per minute for 10 minutes for a total volume of 1 mL. All infusions were calorie matched at 0.5 kcal. Glucose, intralipid and Ensure were dissolved in deionized water. All photometry experiments involving infusions were performed in overnight-fasted AgRP-Cre mice.

To determine whether signaling through GIPR is critical for nutrient-mediated AgRP neuron inhibition, mice equipped for fiber photometry recording and intragastric nutrient infusion were given an injection of a control, non-neutralizing antibody at 30 mg/kg IP^25^ (provided by Eli Lilly) and fasted for 16 hours prior to recordings. At the end of the 16-hour fast, the syringe pump was attached to the intragastric catheter using plastic tubing and adaptors, and mice were habituated to the photometry recording chambers for 20 minutes prior to nutrient infusions. Calorie- and volume-matched infusions of glucose, intralipid, or Ensure were given on different days and recording continued for 10 minutes after the end of infusion. These infusions were completed over the course of 7-10 days, and mice were re-injected with the control antibody every 7 days. After completing these infusions, mice were injected with a previously characterized neutralizing mouse anti-murine GIPR antibody (muGIPR-Ab^25^) at 30 mg/kg and fasted 16 hours before a second round of nutrient infusions was completed as described above. muGIPR-Ab was dosed weekly based on previously published studies^25,29^. AgRP neuron inhibition induced by nutrient infusions was compared across the two conditions. The long-lasting effects of GIPR antibody blockade precluded us from balancing treatment order, and thus recordings following muGIPR-Ab were each performed 7-10 days after control recordings. Additionally, given its long-lasting effects, for all mice that received muGIPR-Ab, subsequent nutrient infusion was a final experiment before euthanasia and confirmation of implant placement and viral expression. To control for possible changes in fiber photometry signal strength over time as a possible cause of muGIPR-Ab effects, a separate cohort of untreated mice were given nutrient infusions at the same time points indicated above without antibody administration.

To determine whether signaling through GLP-1R is critical for nutrient-mediated AgRP neuron inhibition, mice were habituated to the photometry recording chamber for 20 minutes then pre-treated with the GLP-1R antagonist Exendin (9-39) (Ex-9, 1 mg/kg) (HY-P0264, MedChemExpress) or vehicle (saline) 5 minutes prior to infusion of glucose, intralipid or Ensure on separate days. Neural recordings were continued for 10 minutes after the end of infusions. Neural responses to infusions following Ex-9 versus vehicle pretreatment were measured 7-10 days apart for each nutrient, and treatment order was counterbalanced across mice.

### Quantification and statistical analysis

#### Photometry analysis

Photometry data were analyzed with custom Python scripts (https://github.com/nikhayes/fibphoflow), and statistical analyses and data visualizations were generated with Python and Prism. Photometry recordings included emissions from 470nm stimulation and from 405nm stimulation, which were smoothed and downsampled to 1 Hz. Normalization of responses to stimuli relative to baseline was performed on each these signals via the formula: ΔF/F = (F_t_ – F_0_) / F_0_, where F_t_ represents fluorescence at time (t), and F_0_ represents the average fluorescence during the five-minute baseline period preceding the stimulus start time (time zero). To determine statistical significance, the average ΔF/F was calculated for the time frames indicated in the legend for Figures 1, 2, 4, and S1-S5.

#### Behavioral data analysis

To determine chow consumption during fast re-feeding and optogenetic experiments, chow was weighed manually at the specified time points.

#### Statistical analysis

Fiber photometry data were collected and analyzed as previously described^7,26,28^. For photometry traces shown in Figures 1, 2, 4, and S1-S5, ΔF/F (%) refers to the mean ΔF_t_/F_0_*100. For bar graphs quantifying neural responses to chow presentation (Figures 2, S1, S2, and S5), the average ΔF/F over a 1-minute period 3-4 minutes following chow presentation was calculated. For bar graphs quantifying neural responses to nutrient or water infusion (Figures 4, S3, S4, and S5), the average ΔF/F over a 1-minute period at the end of nutrient infusion (9-10min) was calculated. For bar graphs quantifying neural responses to hormone injection (Figures 1, S1, and S2), the average ΔF/F over a 1-minute period 3-4 minutes or 19-20 minutes after injection was calculated as indicated in the figures.

The effects of experimental manipulation versus controls were analyzed with a one-way, repeated-measures ANOVA (Figures 1, 2, S1, S2) or paired T-test (Figures 4, S3, S4, S5) as appropriate for photometry experiments. Fast re-feeding in wildtype mice after treatment with saline or incretin agonists (Figure 2) was analyzed with a one-way, repeated-measures ANOVA.

Food intake in the presence or absence of light stimulation following saline or hormone administration (Figure 3) was analyzed with a 2-way, repeated-measures ANOVA. The Holm-Šídák multiple comparisons test was used as appropriate. Prism was used for all statistical analyses, and significance was defined as p < 0.05. Sample sizes are indicated in the figure legends for each experiment. Where multiple technical replicates of an experiment were performed, trials from the same animal were averaged and handled as a single biological replicate for data analysis and visualization.

